# A CAG repeat threshold for therapeutics targeting somatic instability in Huntington’s disease

**DOI:** 10.1101/2023.12.15.571088

**Authors:** Sarah G. Aldous, Edward J. Smith, Christian Landles, Georgina F. Osborne, Maria Cañibano-Pico, Iulia M. Nita, Jemima Phillips, Yongwei Zhang, Bo Jin, Marissa B. Hirst, Caroline L. Benn, Brian C. Bond, Winfried Edelmann, Jonathan R. Greene, Gillian P. Bates

## Abstract

The Huntington’s disease mutation is a CAG repeat expansion in the huntingtin gene that results in an expanded polyglutamine tract in the huntingtin protein. The CAG repeat is unstable, and expansions of hundreds of CAGs have been detected in Huntington’s disease *post-mortem* brains. The age of disease onset can be predicted partially from the length of the CAG repeat as measured in blood. Onset age is also determined by genetic modifiers, which in six cases involve variation in DNA mismatch repair pathways genes. Knocking-out specific mismatch repair genes in mouse models of Huntington’s disease prevents somatic CAG repeat expansion. Taken together, these results have led to the hypothesis that somatic CAG repeat expansion in Huntington’s disease brains is required for pathogenesis. Therefore, the pathogenic repeat threshold in brain is longer than (CAG)_40_, as measured in blood, and is currently unknown.

The mismatch repair gene *MSH3* has become a major focus for therapeutic development, as unlike other mismatch repair genes, nullizygosity for *MSH3* does not cause malignancies associated with mismatch repair deficiency. Potential treatments targeting *MSH3* currently under development include gene therapy, biologics and small molecules, which will be assessed for efficacy in mouse models of Huntington’s disease. The zQ175 knock-in model carries a mutation of approximately (CAG)_185_ and develops early molecular and pathological phenotypes that have been extensively characterised. Therefore, we crossed the mutant huntingtin allele onto heterozygous and homozygous *Msh3* knock-out backgrounds to determine the maximum benefit of targeting *Msh3* in this model. Ablation of *Msh3* prevented somatic expansion throughout the brain and periphery, and reduction of *Msh3* by 50% decreased the rate of expansion. This had no effect on the deposition of huntingtin aggregation in the nuclei of striatal neurons, nor on the dysregulated striatal transcriptional profile. This contrasts with ablating *Msh3* in knock-in models with shorter CAG repeat expansions. Therefore, further expansion of a (CAG)_185_ repeat in striatal neurons does not accelerate the onset of molecular and neuropathological phenotypes. It is striking that highly expanded CAG repeats of a similar size in humans cause disease onset before 2 years of age, indicating that somatic CAG repeat expansion in the brain is not required for pathogenesis. Given that the trajectory for somatic CAG expansion in the brains of Huntington’s disease mutation carriers is unknown, our study underlines the importance of administering treatments targeting somatic instability as early as possible.

## Introduction

Huntington’s disease is a heritable neurodegenerative disorder, characterised by motor dysfunctions, psychiatric changes, and cognitive decline.^1^ The disease is caused by an expanded CAG repeat in exon 1 of the huntingtin gene (*HTT*), which encodes an expanded polyglutamine (polyQ) tract in the huntingtin protein (HTT). Individuals with a repeat of (CAG)_40_ or more will develop Huntington’s disease in a normal life span, while repeats of around (CAG)_65_ or more will lead to disease onset in childhood or adolescence.^2^ The CAG repeat is unstable in both somatic and germ cells,^2,3^ and large expansions up to approximately (CAG)_1000_ have been observed in patient brains.^4^ In the presence of an expanded CAG repeat, the *HTT* pre-mRNA can be alternatively processed to generate the *HTT1a* transcript, through the activation of two cryptic polyA sites within intron 1.^5,6^ This encodes the highly aggregation prone and pathogenic HTTexon1 protein.^5,7,8^ Huntington’s disease pathology is characterised by the deposition of nuclear and cytoplasmic inclusions,^9,10^ as well as synaptic and neuronal loss in the striatum and other regions of the brain.^11,12^ Nuclear huntingtin aggregation is linked to transcriptional dysregulation, a molecular phenotype observed in mouse models and *post-mortem* brains.^13–15^

Although the age of disease onset is inversely correlated with CAG repeat length, genetic and environmental modifiers also contribute to the variation in age of onset and disease progression.^16^ Genome-wide association studies (GWAS) have identified potential genetic modifiers, a subset of which have variants associated with genes involved in DNA damage repair pathways including *FAN1*, *MSH3*, *MLH1*, *LIG1*, *PMS1* and *PMS2*.^17–20^ That the variants affected the function of these DNA repair genes was supported by previous experiments showing that the ablation of specific mis-match repair genes (*Msh2*, *Msh3*, *Mlh1* and *Mlh3*) reduced somatic CAG repeat instability in mouse models of Huntington’s disease.^21–23^ Mechanistically, it has since been shown that FAN1 nuclease activity is implicated in slowing CAG repeat expansion,^24^ that the promotion of somatic CAG instability by FAN1 is blocked in the absence of MLH1^25^ and that FAN1 controls mismatch repair assembly via MLH1 to stabilise the CAG repeat.^26,27^

It is now widely accepted that the somatic expansion of a mutant CAG repeat in brain cells is the first step in the molecular pathogenesis of Huntington’s disease, followed by a second process that leads to cell dysfunction, once a CAG repeat threshold has been achieved. Therefore, it is the rate of somatic CAG repeat expansion that drives the age of onset and rate of progression of the disease, and the genetic modifiers that impact this process are potential therapeutic targets.^28,29^ Of these, MSH3, part of the MutSβ heterodimer that is thought to recognise the potential slip outs caused by the expanded CAG repeats,^30,31^ is of particular interest. Previous work has established that *Msh3* is required for somatic instability in the *Hdh*Q111 mouse model of Huntington’s disease,^32^ and that variants of mouse *Msh3*^33^ and levels of *MSH3* expression in Huntington’s disease patients^34^ affect somatic expansion. Importantly, for therapeutic potential, unlike other DNA mis-match repair genes, loss of function of MSH3 does not cause Lynch syndrome, which is associated with a high risk of colon cancer and a spectrum of other malignancies.^35,36^

The design of mouse trials for the target validation of MSH3 and for the preclinical testing of novel MSH3 targeting therapeutics is not straightforward. Ideally, the chosen model will develop robust phenotypes within a timescale amenable to the administration and durability of the potential therapeutic agents. However, such a model is likely to have been created with a highly expanded CAG repeat that would cause childhood onset of the disease in humans, and therefore, does not model the CAG repeat expansion process that occurs in the adult-onset disease. The zQ175 knock-in mouse of Huntington’s disease is an attractive candidate as it is well characterised^7,37,38^ and has been used for HTT lowering approaches.^39^ However, the mutant allele in zQ175 generally has approximately (CAG)_190_. To test whether the prevention of somatic CAG repeat expansion could result in measurable improvements in zQ175 mice, we crossed the mutant *Htt* allele onto heterozygous and homozygous *Msh3* knockout backgrounds. Although, in the absence of *Msh3*, the pronounced somatic CAG repeat instability in the striatum was prevented, there was no improvement in striatal transcription dysregulation, as measured by RNA-seq, nor was there a decrease in striatal HTT aggregation. These data point to a CAG repeat threshold above which the rate of disease pathogenesis does not increase, and at which the prevention of somatic CAG repeat expansion will have no beneficial consequences. The trajectory for CAG repeat-expansion in the human brain is not known and this work further underlines the importance of developing treatments that can be given to Huntington’s disease mutation carriers as early as possible.

## Materials and methods

### Ethics statement

All procedures were performed in accordance with the Animals (Scientific Procedures) Act, 1986, and approved by the University College London Ethical Review Process Committee.

### Generation of *Msh3* knock out mice

*Msh3* knockout mice were generated by CRISPR technology.^40^ In brief, guide RNAs (gRNAs) targeting exon 3 of mouse *Msh3*: Msh3 gRNA 51/57 (targeting sequence: TCTGCAACGTGCAAAGAATGCGG) and exon 8, Msh3 gRNA 89/69, (Targeting sequence: GTTGGCACAGACAGGTCGGAGGG) were designed by Benchling online software and generated by *in vitro* transcription. Fertilised eggs from C57BL/6J mice were injected with the two gRNAs (Msh3 gRNA 51/57 and Msh3 gRNA 89/69) and Cas9 protein (PNA BIO INC) and transferred to pseudopregnant CD1 female mice. *Msh3*^+/−^ founders (F0) mice were backcrossed to C57BL/6J (Charles River) mice and maintained on C57BL/6J background (Charles River).

### Animal colony breeding and maintenance

zQ175 mice on a C57BL/6J background were obtained from the CHDI Foundation colony at the Jackson Laboratory (Bar Harbour, Maine). zQ175 males were bred to *Msh3*^+/−^ females to generate zQ175:*Msh3*^+/−^ males that were then bred to *Msh3*^+/−^ females to generate the six required genotypes as littermates and aged to 6 months. The mean and standard deviation of the CAG repeat sizes were zQ175 186 ± 2.3, zQ175:*Msh3^+/−^* 185 ± 2.4 and zQ175:*Msh3*^−/−^ 182 ± 2.0. A second smaller generation was bred and aged to 3 months, with CAG sizes zQ175 185 ± 3.2, zQ175:*Msh3^+/−^* 183 ± 2.6 and zQ175:*Msh3*^−/−^ 190 ±6.0. The CAG repeat size for the 2-month-old zQ175 mice for HTRF was 192 ± 6.4.

Mouse husbandry and health monitoring were carried out as previously described.^41^ Mice were group-housed by gender, with mixed genotypes per cages. Animals were kept in individually ventilated cages with Aspen Chips 4 Premium bedding (Datesand) and environmental enrichment through access to chew sticks and a play tunnel (Datesand). All mice had constant access to water and food (Teklad global 18% protein diet, Envigo, the Netherlands). Temperature was regulated at 21°C ± 1°C and mice were kept on a 12 h light/dark cycle. The facility was barrier-maintained and quarterly non-sacrificial FELASA (Federation of European Laboratory Animal Science Associations) screens found no evidence of pathogens. Tissues were harvested, snap frozen in liquid nitrogen and stored at −80°C. Mice were transcardially perfused with 4% paraformaldehyde as previously described.^7^

### Real-time quantitative PCR for *Msh3* transcripts

RNA was isolated and real-time quantitative PCR (qPCR) was performed as previously described.^42^ Taqman assays to the deleted *Msh3* region (Thermo, Mm00487756_m1) and outside the deleted region (Thermo, Mm01290051_m1) were used, and to housekeeping genes for *Atp5b* and *Eif4α2* as described previously.^42^

### Genotyping and CAG repeat sizing

Genomic DNA isolation from ear biopsies or from frozen brain regions and peripheral tissues was carried out as previously described.^43^ DNA from ear biopsies was diluted to 50 ng/μL. The DNA concentration from brain regions and peripheral tissues was variable but within the range of 100 – 300 ng/μL.

Genotyping PCRs for zQ175 mice were performed as previously described.^43^ The *Msh3* PCR was carried out in 10 μL reactions using DreamTaq Buffer (Thermo). The *Msh^−/−^* knockout allele was amplified with primers Msh3 51 SF (5′-GAATTGTCCTAGTGAAGCCAAAA-3′) and Msh3 89SR (5′-CTTGGGAGGAAAAGCAGATG-3′) generating a product of 501 bp. The wild-type allele was amplified with primers Msh3 51 SF and Msh3 51 SR (5′-AACCAACTACTCTCCCCCAAA-3′) generating a 479 bp product. Primers were at 5 pmol/μL, and cycling conditions were 2.5 min at 95 °C, 32 x (20 s at 95 °C, 30 s at 56 °C, 1 min at 72 °C), 6 min at 72 °C.

CAG repeat sizing for ear biopsy DNA was carried out as described previously and for somatic instability analysis of DNA from tissue samples, reaction volumes were scaled to 25 μL.^43^ The PCR product was denatured in HiDi Formamide (Thermo Fisher Scientific) and run in triplicate on ABI3730XL Genetic Analyser with MapMarker ROX 1000 (Bioventures) internal size standards and analysed using GeneMapper v6 software (Thermo Fisher Scientific). The tissue-wide analysis of mode and instability index was analysed as described,^27,44^ using a custom R script.^27^ A more detailed statistical analysis of striatal samples was run using JMP^®^, v17 (SAS Institute Inc., Cary, NC).

### Western blotting

Cortex was lysed on ice in KCL buffer (50 mM Tris pH 8.0, 10% Glycerol, 150 mM KCL, 5 mM EDTA, with 1 mM phenylmethylsulphonyl fluoride and cOmplete protease inhibitor cocktail tablets (Roche)). Tissue homogenisation and western blotting was performed with 50 μg of total protein as previously described.^15^ Membranes were incubated overnight with gentle agitation at 4 °C with the MSH3 antibody (Merck MSH3MAB(1F6)) in 5% milk PSB-T (phosphate buffer saline (PBS), 0.1% Tween-20). Tubulin was detected by incubating for 1 h at room temperature in 5% Milk-PBST (Marvel) with an α-tubulin antibody (Sigma T9026) at a 1:40.000 dilution.

### Tissue lysis and QuantiGene assays

The lysis of tissue samples and the QuantiGene multiplex assays were carried out as previously described,^42^ with 1:2 dilutions for each brain region. All QuantiGene probe sets are available from Thermo Fisher Scientific and the individual accession numbers and probe set regions are summarised in Supplementary Figure 1 and Supplementary Tables 1 and 2. The sequences for the capture extenders, label extenders and blocking probes are available from Thermo Fisher Scientific on request. Data analysis was performed by normalising raw MFI signals obtained to the geometric mean of the housekeeping gene signals.

### Homogeneous time-resolved fluorescence

Homogeneous time-resolved fluorescence (HTRF) was performed as previously described,^7^ using *n* = 8/ genotype/ age with an equal ratio of males to females. The antibody pairings and concentrations for the soluble HTTexon1 assay were donor: 2B7-Tb, 1 ng and acceptor: MW8-d2, 40 ng. For the aggregated HTTexon1 assay the antibody pairings were donor: 4C9-Tb, 1 ng and acceptor: MW8-d2, 10 ng. The donor and acceptor antibodies were added in 5 μL HTRF detection buffer (50 mM NaH_2_PO_4_, 0.2 M KF, 0.1% bovine serum albumin, 0.05% Tween-20) to give the optimised antibody concentration when added to 10 μL of a 10 % lysate concentration in a 384-well ProxiPlate (Greiner Bio-One). Following incubation for 3 h on an orbital shaker (250 rpm), plates were read using an EnVision (Perkin Elmer) plate reader as previously described.^43^

### Immunohistochemistry

The immunostaining and imaging of coronal sections using S830 was carried out as previously outlined.^7^ Sections from 3- and 6-month-old mice were stained and imaged separately, but the four genotypes for each age were processed together.

To perform an analysis of aggregation throughout the striatum, four images per striatum across five hemispheres/sections were captured (20 images in total). Images were obtained using a Zeiss AxioSkop2 Plus microscope fitted with a Zeiss AxioCam 208 colour camera and were recorded using Zeiss ZEN 3.2 (blue edition). For quantification and thresholding, images were converted to 8-bit in ImageJ and a signal intensity level was chosen to indicate staining above background. Signals were segmented based on pixel intensity and the number of positive pixels in contact with each other. A gate of 80 pixels was applied to all signals over 190 to identify S830 positive objects. Inclusions were identified by applying a gate of 2 pixels and a signal of 150, the vast majority of which would be nuclear inclusions. The average area of each object and the percentage of the image that contained S830 immunostaining was recorded.

### RNA Sequencing

Tissues were homogenised in Qiazol lysis reagent (Qiagen) and RNA was extracted from the same set of samples as used for somatic instability. Total RNA isolation was performed as outlined^42^ using the RNeasy mini kit (Qiagen). RNA concentration (Thermo, Q32855), integrity and quality (Thermo, Q33222) were measured using a Qubit Fluorometer (Themo Fisher Scientific Q33238). RNA sequencing and analysis was then performed as described previously.^45^ RNA was converted into cDNA, indexed and libraries sequenced on the Illumina NovaSeq 6000 system to generate 30 million paired-end reads. Differential expression tests were performed in R using DESeq2^46^ with independent filtering disabled, and a significance threshold of an adjusted *P* < 0.05 after multiple-test correction, and fold changes of at least 20% in either direction. Analysis of RNA-Seq data comparing zQ175 with wild-type to generate the “HD signature” lists of transcriptionally dysregulation genes was also based on the same significance criteria. The posterior probabilities-based method^45^ was used to quantify the probability that *Msh3* knockout out restored transcriptional dysregulation, either partially or fully, on a gene-by-gene basis. This was done by viewing the effect of *Msh3* genotype as a multiple of the HD model signature (in log2). If α < 0, then the disease effect is reversed by *Msh3* knockout, if α > 0, the disease effect is exacerbated by loss of *Msh3*. The thresholds for each type of reversal class were as previously calculated^45^ and were as follows: super-reversal: α < –1.3; full reversal: –1.3 < α < –0.7; partial reversal: –0.7 < α < –0.3; negligible reversal: –0.3 < α < 0.3; and exacerbation: α > 0.3. The genes were then scored for normalization based on the overall reversal probability. The ‘HD signature’ dysregulated gene list was tested for gene set overrepresentation against the GOBP (gene ontology biological process)^47^ version 18.9.29 using the enricher function within the R clusterProfiler^48^ package. A clusterProfiler *q* < 0.05 was deemed significant.

### Statistical analysis

Data were screened for outliers at sample level using the ROUT test (Q = 10%; GraphPad Prism v9) and outliers were removed from the analysis. All data sets were tested for a normal gaussian distribution (Shapiro-wilk, Prism v9). Statistical analysis was by one-way ANOVA, two-way ANOVA or mixed-effects model (REML) with either Tukey’s or Bonferroni *post hoc* tests. Non-parametric analysis was performed by the Kruskal-Wallis test with Dunn’s *post hoc* analysis. Graphs were prepared using Prism v9 (GraphPad Software, California, USA). *P*-values <0.05 were considered statistically significant. PCA analysis of the summary statistics for the CAG repeat profiles of the striatal and ear tissue was performed. The Standard Least Squares model with Effect Leverage was conducted with ear trace data included as a covariate. Data for variance were log transformed and graphs prepared using JMP^®^ v17 (SAS Institute Inc., Cary, NC).

## Results

An *Msh3* knockout mouse line was generated using CRISPR Cas9 editing,^40^ targeting exons three to eight as shown in Fig. 1A. A colony was established on a C57BL/6J background and mice were bred to homozygosity. The level of *Msh3* mRNA in cortical lysates was shown to have decreased by 50% in heterozygous (*Msh3*+/−), and to have been ablated in homozygous (*Msh3-/-*), knockout mice by real-time quantitative PCR (qPCR) (Fig. 1B). MSH3 protein levels were determined in cortical lysates by western blotting and similarly showed that MSH3 had been reduced to at least 50% in *Msh3*+/− heterozygotes and could not be detected in the *Msh3*-/- homozygotes (Fig. 1C). To determine the effects of MSH3 reduction and MSH3 ablation in zQ175 mice, zQ175 males were bred to *Msh3+/−* females to generate zQ175 males that were heterozygous for *Msh3* knockout. These were then bred to *Msh3+/−* females to generate all six genotypes of interest as littermates in a single generation (Fig. 1D).

**Figure 1.**
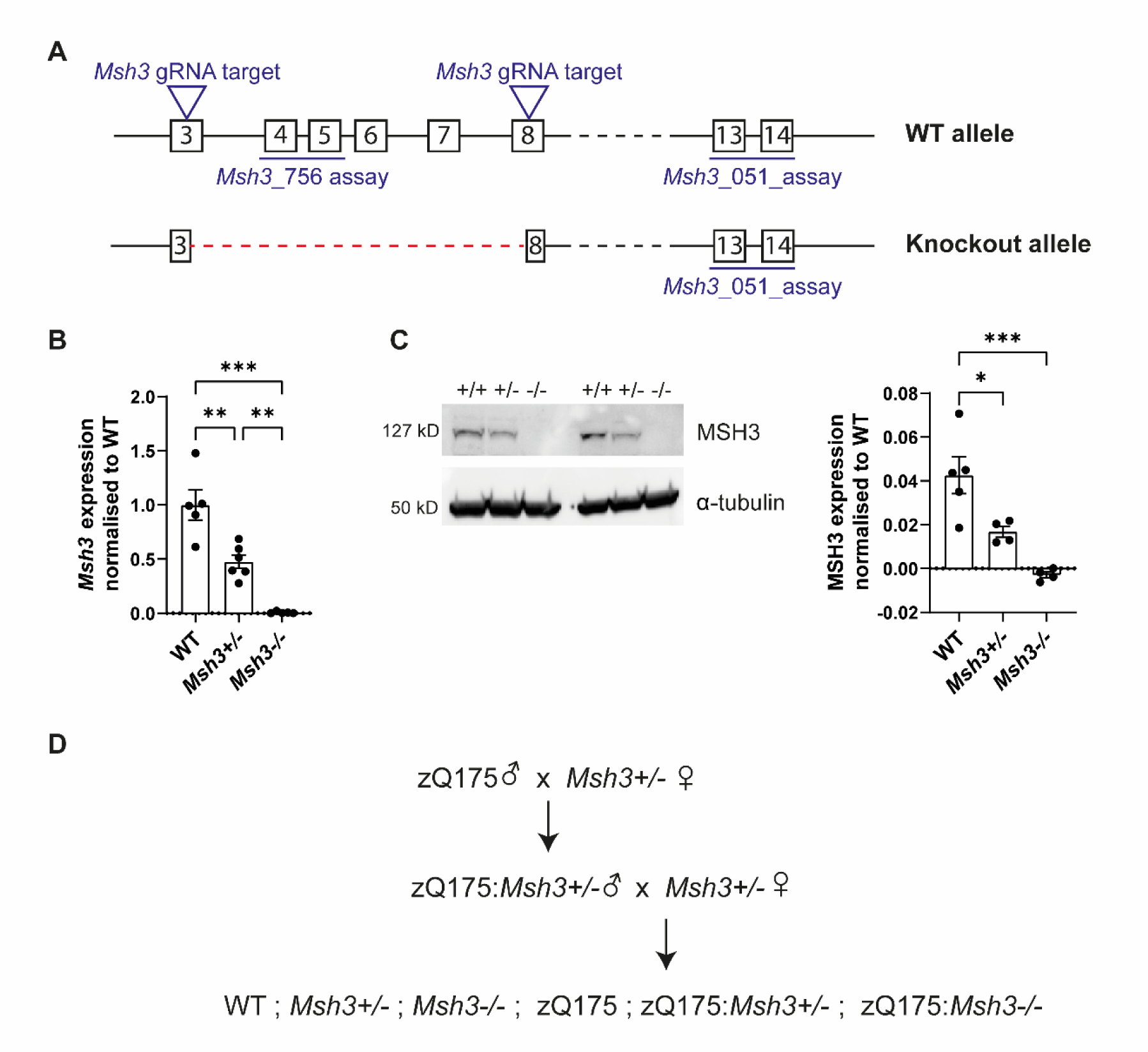
Characterisation of the *Msh3* knockout line. **(A)** *Msh3* knockout allele generation by CRISPR/Cas9 showing the position of guide RNAs (gRNA) targeting *Msh3* exons 3 and 8 and the position of the qPCR assays. **(B)** Expression of the *Msh3* mRNA was confirmed, using a qPCR assay with primers in the deleted region (*Msh3*_756), in the cortex of heterozygous *Msh3*+/− and homozygous *Msh3*-/- mice at 8 weeks of age (*N* = 5/genotype). **(C)** Western blotting with an MSH3 antibody showed that MSH3 levels were decreased in *Msh3*+/− heterozygotes and absent in *Msh3*-/- homozygotes. MSH3 levels were quantified by normalising to α-tubulin (*N* = 5/wild-type, N = 4/*Msh3*+/− and *Msh3*-/-). **(D)** Schematic of the genetic cross performed to generate the six experimental genotypes as littermates. One-way ANOVA with Tukey’s post hoc test. Error bars = SEM. \**p* ≤ 0.05, \*\**p* ≤ 0.01, \*\*\**p* ≤ 0.001. The test statistic, degrees of freedom and *p* values are summarised in Supplementary Table 3. Full-length blots are shown in Supplementary Fig. 5. WT= wild-type.

### Loss of *Msh3* prevents somatic CAG repeat instability in zQ175 mice

We first investigated the extent of somatic CAG repeat instability in ten CNS regions and several peripheral tissues from zQ175 mice at 6 months of age (N = 8). The GeneMapper CAG repeat profiles for the brain regions and tissues for each mouse were compared to that obtained from the ear sample taken at post-natal day 12 (P12) (Fig. 2). Within the CNS, instability was most pronounced in the striatum, brain stem, colliculus, thalamus, hypothalamus and spinal cord, consistent with that observed in other mouse models.^49^ In the periphery, instability was extensive in the liver as expected,^23,33,49^ but absent from the heart. Unexpectedly, CAG repeat contraction had led to a reduction in the CAG repeat length mode in the pituitary. Examination of the traces for zQ175:*Msh3*-/- mice showed that nullizygosity for *Msh3* prevented CAG repeat expansion and that the repeat contractions had resulted in a decrease in the mode in most brain regions and peripheral tissues (Fig. 2). Heterozygosity for *Msh3* in zQ175:*Msh3*+/− mice caused in a reduction in the extent of repeat expansion (Fig. 2).

**Figure 2.**
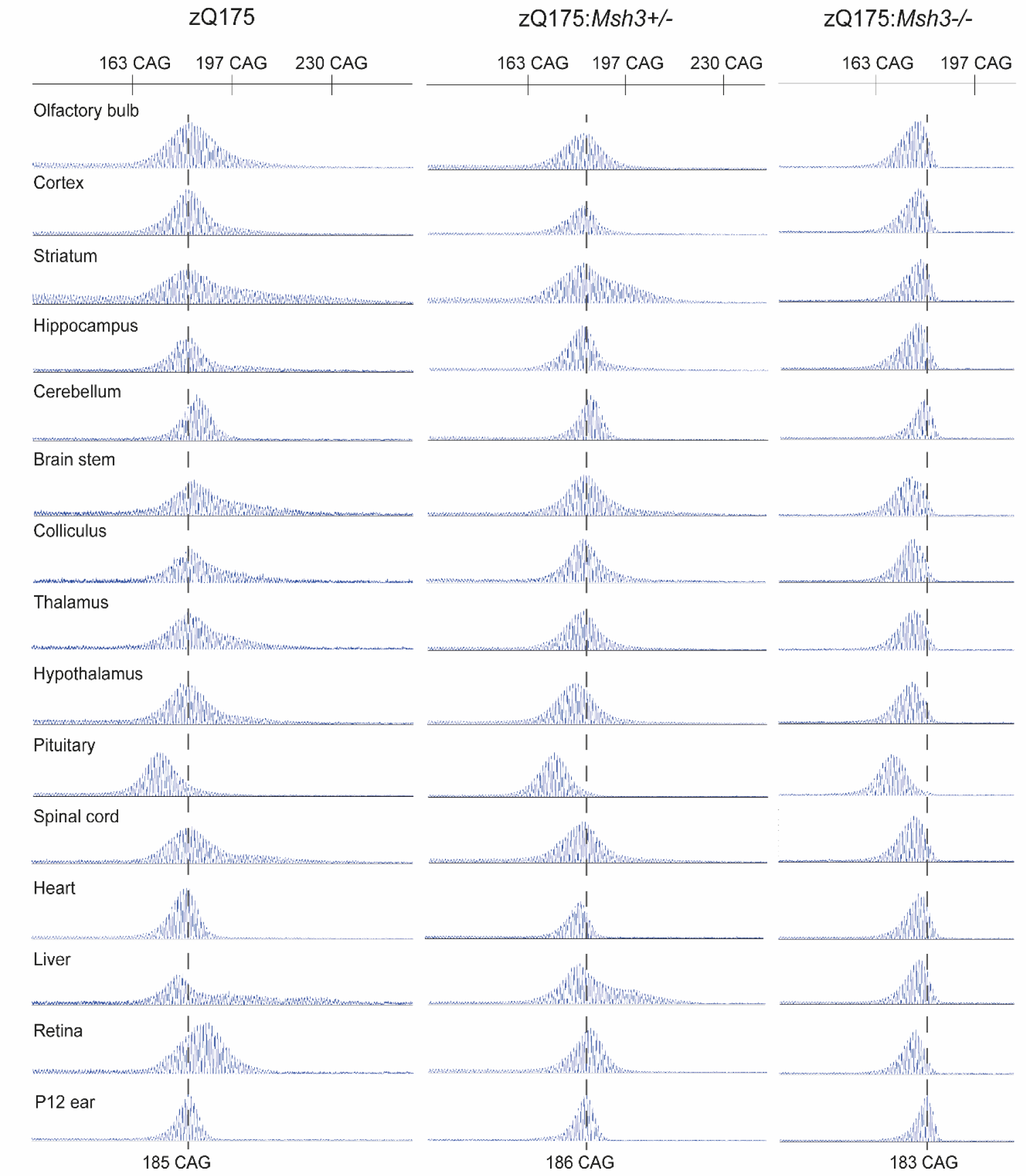
CAG repeat instability in zQ175 mice is dependent on *Msh3* expression. Representative GeneMapper traces of CAG repeat profiles for brain regions and peripheral tissues from mice at 6 months of age (*N* = 8/genotype). The modal CAG repeat length in ear samples at 12 days of age (P12) for zQ175 mice was 186 ± 2.3, for the zQ175:*Msh3+/−* mice was 185 ± 2.4 and for zQ175:*Msh3*-/- mice was 182 ± 2.0. The representative traces are from the same mouse for each genotype with the CAG repeat mode at P12 indicated by the dashed line. Somatic instability in 6-month-old zQ175 mice was most pronounced in the striatum, brainstem, spinal cord, and liver, and the repeat had contracted in the pituitary. Somatic CAG expansion was prevented in zQ175 mice in which *Msh3* was absent, and contractions had led to a reduction in the CAG repeat mode. Heterozygosity for *Msh3* led to a decrease in the extent of repeat expansion.

The instability index quantifies the change in CAG repeat length profiles between samples.^44^ This was calculated for each brain region and peripheral tissue for every mouse by comparing its CAG repeat trace with that of the P12 ear sample (N = 8/genotype). Plotting the change in the mean instability index for each tissue, demonstrated the highly significant, stepwise reduction in CAG repeat instability with loss of MSH3 (Fig. 3A). In many of the brain regions and tissues, e.g. olfactory bulb, colliculus thalamus and retina, the level of instability in zQ175:*Msh3*+/− mice was decreased to approximately 50% of the reduction observed in the zQ175*Msh*3-/- mice. In some brain regions, e.g. striatum, the decrease in instability index between zQ175 and zQ175:*Msh3*+/− mice was less pronounced than between zQ175:*Msh3*+/− and zQ175:*Msh3*-/- mice.

**Figure 3.**
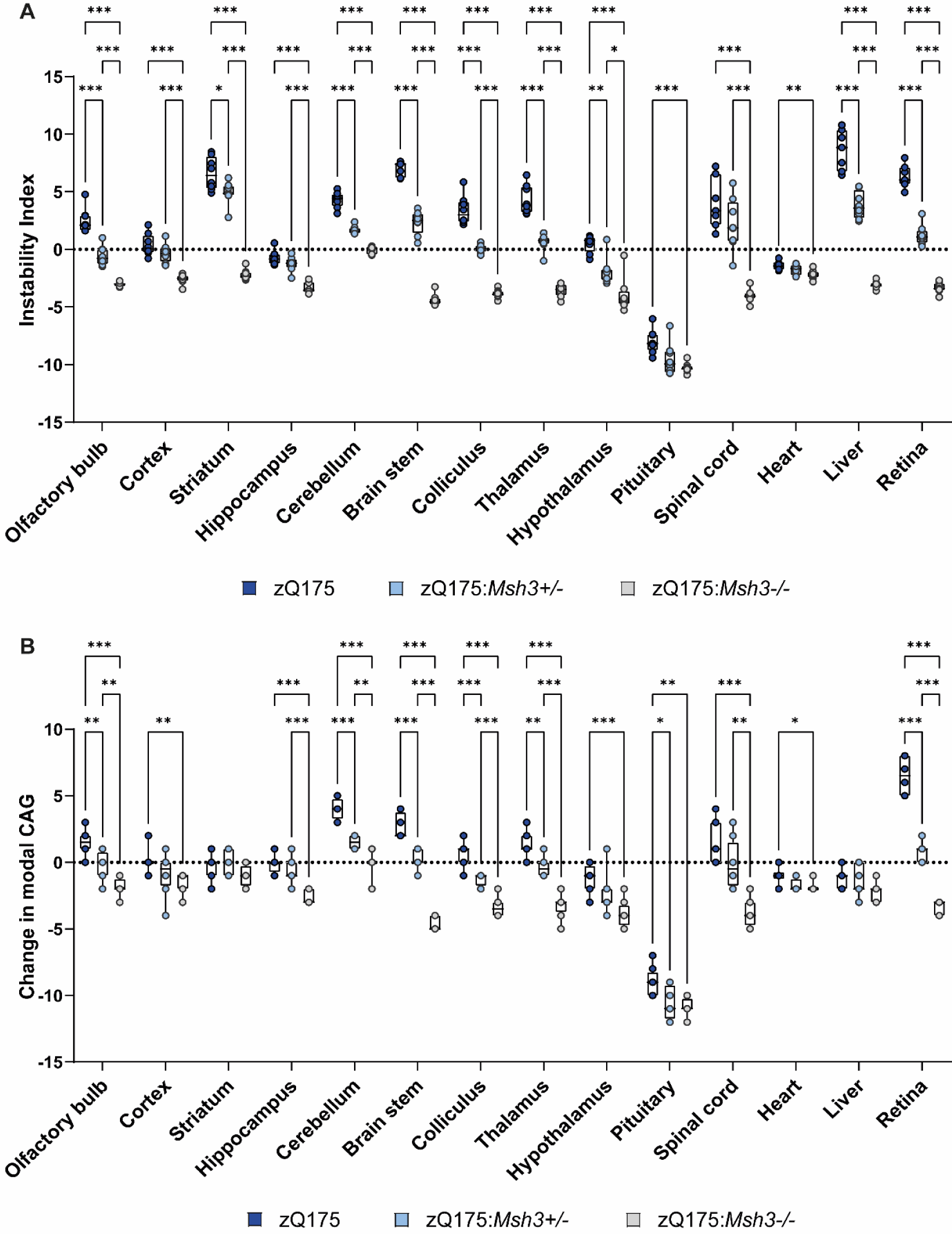
Tissue specific changes in the instability index and CAG repeat mode of the of the CAG repeat profiles in zQ175, zQ175*:Msh3*^+/−^ and zQ175:*Msh3*^−/−^ mice. **(A)** Instability index quantified for the GeneMapper traces for each brain region and tissue normalised to P12 ear sample for each genotype. A stepwise reduction in the instability index was observed for each brain region / tissue with the reduction and loss in *Msh3* expression. **(B)** Quantification of the change in mode from the P12 ear sample. Modal CAG was reduced with *Msh3* genotype, in all regions apart from striatum, heart and liver*. N* = 8/genotype, 2-3 technical replicates for each mouse. \**p* ≤ 0.05, \*\**p* ≤ 0.01, \*\*\**p* ≤ 0.001. Box and whisker plots represent median and 25^th^ and 75^th^ percentiles with bars drawn from minimum to maximum. Statistical analysis was mixed-effects model (REML), with Tukey’s correction for multiple comparisons. The test statistic, degrees of freedom and *p* values are summarised in Supplementary Table 4.

The change in mode CAG repeat length between each brain region and tissue at 6 months of age as compared to the P12 ear sample was also calculated for each mouse (N = 8/genotype). The average mode CAG length changed little with age in most brain regions and tissues for zQ175 mice, exceptions being the cerebellum, brain stem, pituitary, and retina (Fig. 3B). However, there was a reduction in the average mode CAG length in the zQ175:*Msh3-*/- mice in most brain regions and tissues (Fig. 3B).

As the most somatically unstable region, we set out to analyse the striatal traces in more detail using summary statistics in a multivariate analysis. The Principal Component Analysis (PCA) investigated correlations between a set of summary statistics of CAG repeat profiles and found new independent trends (Fig. 4A). The first two components explained 77.3 % of the total variation with the first component explaining 62.2 % alone. The first component in the scores plot separated the striatum for the zQ175:*Msh3*+/− and zQ175:*Msh3*-/- mice from the ear tissues of all three genotypes and the striatum of the zQ175 mice. To a lesser degree the second component separated the ear tissue from the striatum of the zQ175 mice. The loadings plot in Fig. 4B represents the summary statistics, and the relationship between them. The summary statistics related to location (mean/median/mode) and variation (log variance/log interquartile range (IQR) along with skewness, instability and expansion were all closely correlated in the first component, with contraction and kurtosis negatively correlated in the second component. The information from both the scores and loadings plots confirmed the difference seen visually with the traces in Fig 2, and that these can be attributed to specific summary statistic differences.

**Figure 4.**
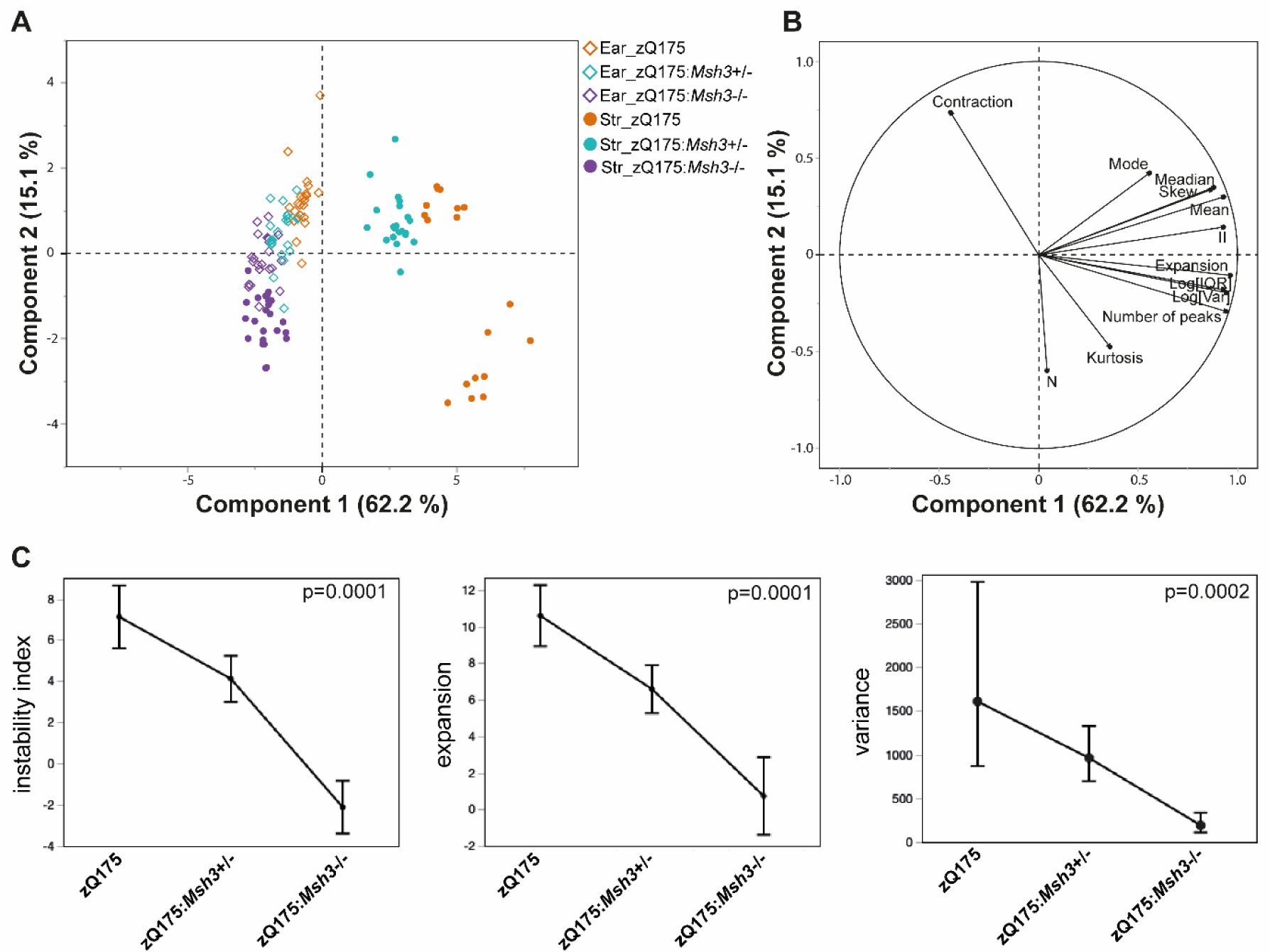
Statistical analysis of the effect of heterozygosity and homozygosity for *Msh3* knock-out on somatic CAG repeat instability in the striatum. **(A-B)** Scores and Loading plots from the PCA of all summary statistics using ear and striatal data. The plots indicate the statistical variables contributing to the differences seen between genotypes: contraction, mode, median, skew, mean, expansion, II = instability index, log(IQR) = log interquartile range, log(var) = log variance, kurtosis and N = sum of peak heights. **(C)** Least Squares Means plots of instability index, expansion, and variance summary statistics showing means (geometric means for variance) adjusted for ear and 95% confidence intervals for striatum across genotype. These data again confirm the significant effect of *Msh3* genotype on instability, expansion, and variation. *P*-value of effect test (genotype) shown.

We also analysed specific variables of the striatal traces in more detail; instability index, expansion, and variation (Fig. 4C). The instability index as calculated in Fig 3. A is normalised to the ear as a control, however due to the genetic nature of the intervention, these ear samples were also affected by the genotype, as visible in the zQ175:*Msh3*-/- ear trace in Fig 2. We reasoned that instability normalised to the ear trace, whilst a good indication of expansion, may not be the most accurate, as the resulting ratio could have an inflated variability. Therefore, we used a standard least squares model, and included the ear data of each variable, as a co-variate. This analysis of covariance (ANCOVA) approach enabled us to ensure that any difference detected indicated a true effect of the genotype on the striatum, over and above the non-specific effect. These results again demonstrated a clear effect of genotype on the instability index, expansion, and variation of the striatal traces, confirming the effect of *Msh3* genotype on instability, and other statistical variables that characterise the traces seen in Fig 2.

### The absence of MSH3 had no effect on transcriptional dysregulation in the striatum of zQ175 mice

Transcriptional dysregulation is a molecular phenotype that has been well-characterised in Huntington’s disease *post-mortem* brains and mouse models^14,50^. This is particularly the case for the striatal transcriptome of an allelic series of knock-in mice^13^ for which a robust disease signature has been established.^51^ Therefore, we performed RNA-seq on striata from wild-type, zQ175, *Msh3*-/- and zQ175:*Msh3*-/- mice to investigate the effect of *Msh3* ablation on the striatal transcriptome of zQ175 mice at 6 months of age.

We began by comparing the expression levels of striatal genes between zQ175 and wild-type mice. The extent of transcriptional dysregulation is illustrated in the volcano plot (Fig. 5A). This zQ175 signature correlated with those previously obtained for the striata from the zQ175, Q140 and *Hdh*Q111 knock-in mouse models at 6 and 10 months of age (Supplementary Fig. 2A).^13^ Similarly, the gene set overrepresentation of the 2,486 dysregulated genes allocated to gene ontology biological process (GOBP) confirmed that the transcriptional dysregulation in the zQ715 mice was consistent with that previously reported (Supplementary Fig. 2B).^45^

**Figure 5.**
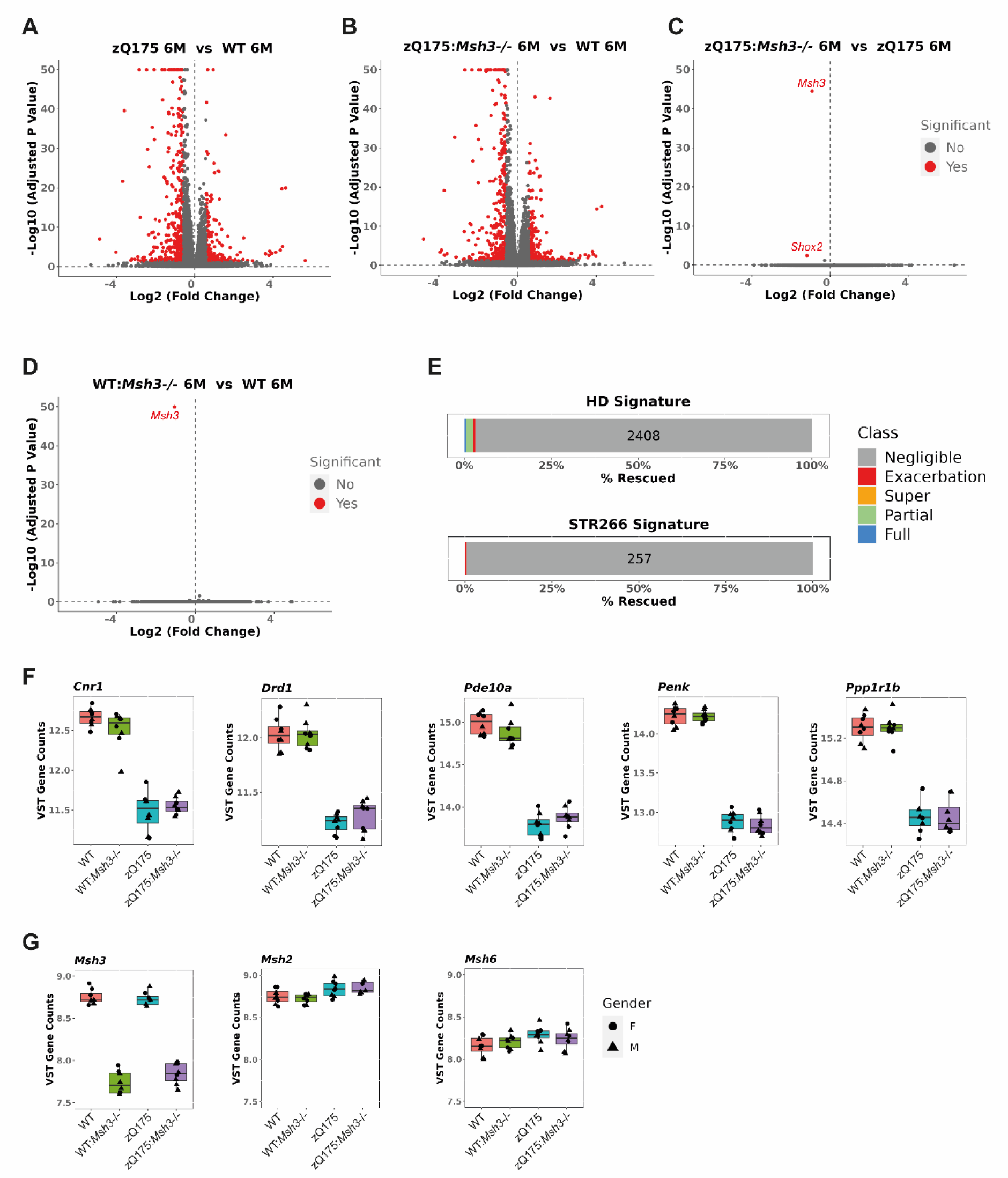
Transcriptional dysregulation in the striatum of zQ175 mice is not alleviated by loss of *Msh3* expression. (A-D) Volcano plots illustrating gene expression changes between. **(A)** zQ175 and wild-type mice **(B)** zQ175:*Msh3*-/- and wild-type mice, **(C)** zQ175 and zQ175:*Msh3*-/- mice and **(D)** *Msh3*-/- and wild-type mice, at 6 months of age. **(E)** Of the 2486 genes dysregulated in zQ175 mice as compared to WT mice at 6 months of age, 2408 showed negligible reversal, with a total of 15 genes showing full reversal, and 51 partial reversal, indicating that only 3% of genes were rescued, a level considered to be background. Of the known 266 genes dysregulated in striatal tissues across models, 258 were detected as dysregulated, with none reversed and 1 exacerbated. **(F-H)** Expression levels for **(F)** striatal genes of interest: *Cnr1, Drd1, Pde10a, Penk,* and *Ppp1r1b* and **(G)** the DNA mismatch repair genes *Msh3, Msh2, Msh6* plotted as variance stabilizing transformed counts derived using DESeq2. Box and whisker plots represent median and 25^th^ and 75^th^ percentiles with bars drawn from minimum to maximum. They show there was no change in expression due to loss of *Msh3*. *N* = 8/genotype, 4 males/4 females for each genotype except zQ175*:Msh3*-/- at 5 males/3 females. WT= wild-type.

The extent of transcriptional dysregulation in both zQ175 and zQ175:*Msh3*^−/−^ mice as compared to wild-type was very similar (Fig. 5B). Comparison of the striatal gene expression profiles between zQ175 and zQ175:*Msh3*-/- identified only two changes, *Msh3* and *Shox2*, the most significant of which was *Msh3* (Fig. 5C). The ablation of MSH3 resulted in the expression of only 3% of the 2,486 genes dysregulated in the zQ175 striata being reversed toward wild-type levels whilst the dysregulated levels of 1% of these genes was exacerbated (Fig. 5E). Of the 65 genes for which the expression level was reversed, 14 were fully reversed and 51 partially reversed, numbers corresponding to below background (Fig. 5E). We also considered the 266 genes that comprise the ‘striatal signature’ of robustly dysregulated genes across multiple Huntington’s disease models.^51^ Of these, 258 were dysregulated in the zQ175 striatum, and a reversal of expression toward wild-type levels in the absence of MSH3 was not observed for any of these genes (Fig. 5E). The expression levels of a few key striatal genes in wild-type, Msh3-/-, zQ175 and zQ175:Msh3-/- striata are illustrated in Fig. 5F.

Ablation of MSH3 had no effect on the striatal transcriptome of wild-type mice; *Msh3* being the only transcript found to differ between the two genotypes (Fig. 5D). Consistent with this, ablation of MSH3 had no effect on the expression level of a panel of mis-match repair genes in either wild-type or zQ175 mice in the striatum (Fig. 5G and Supplementary Fig. 3) or other brain regions (Supplementary Fig. 3) as measured by a QuantiGene multiplex assay.

### Striatal nuclear HTT aggregation levels were unchanged with loss of *Msh3* in zQ175 mice

Several lines of evidence indicate that the retention of aggregated HTT in neuronal nuclei precedes transcriptional dysregulation.^7,13,15^ Therefore, we used immunohistochemistry with the S830 anti-HTT antibody to quantify the level of HTT aggregation in striata. Using this approach, we have previously shown that aggregated HTT could first be detected in striatal nuclei from zQ175 mice before two months of age as a diffuse immunostain that filled the nuclei.^7^ By six months of age, this had often resolved to a nuclear inclusion either with or without the diffusely distributed aggregated HTT.^7^ Coronal sections from zQ175, zQ175:*Msh3*^+/−^ and zQ175:*Msh3*-/- mice, together with wild-type controls, at three and six months of age were immunostained with S830. At three months of age, the S830 immunostain appeared diffusely nuclear as expected, with the initiation of inclusion bodies being detected in some cells (Fig. 6A). The application of thresholding and quantification of signal intensity indicated that there was no difference in the number of S830 positive objects, the average area of these objects and the percentage of the image stained between the three genotypes (Fig. 6B). Therefore, *Msh3* genotype did not influence the first appearance of HTT aggregation in striatal nuclei. At six months of age, in addition to the diffuse HTT aggregation, nuclear inclusions were more prominent (Fig. 6C). Thresholding was applied to quantify the number of S830 positive objects, their average size and the percentage of the image stained both for all objects (Fig. 6D) and for inclusions, the majority of which were nuclear (Fig. 6E). As at three months of age, at six months, there was no difference in the level of HTT aggregation in striatal nuclei between genotypes.

**Figure 6.**
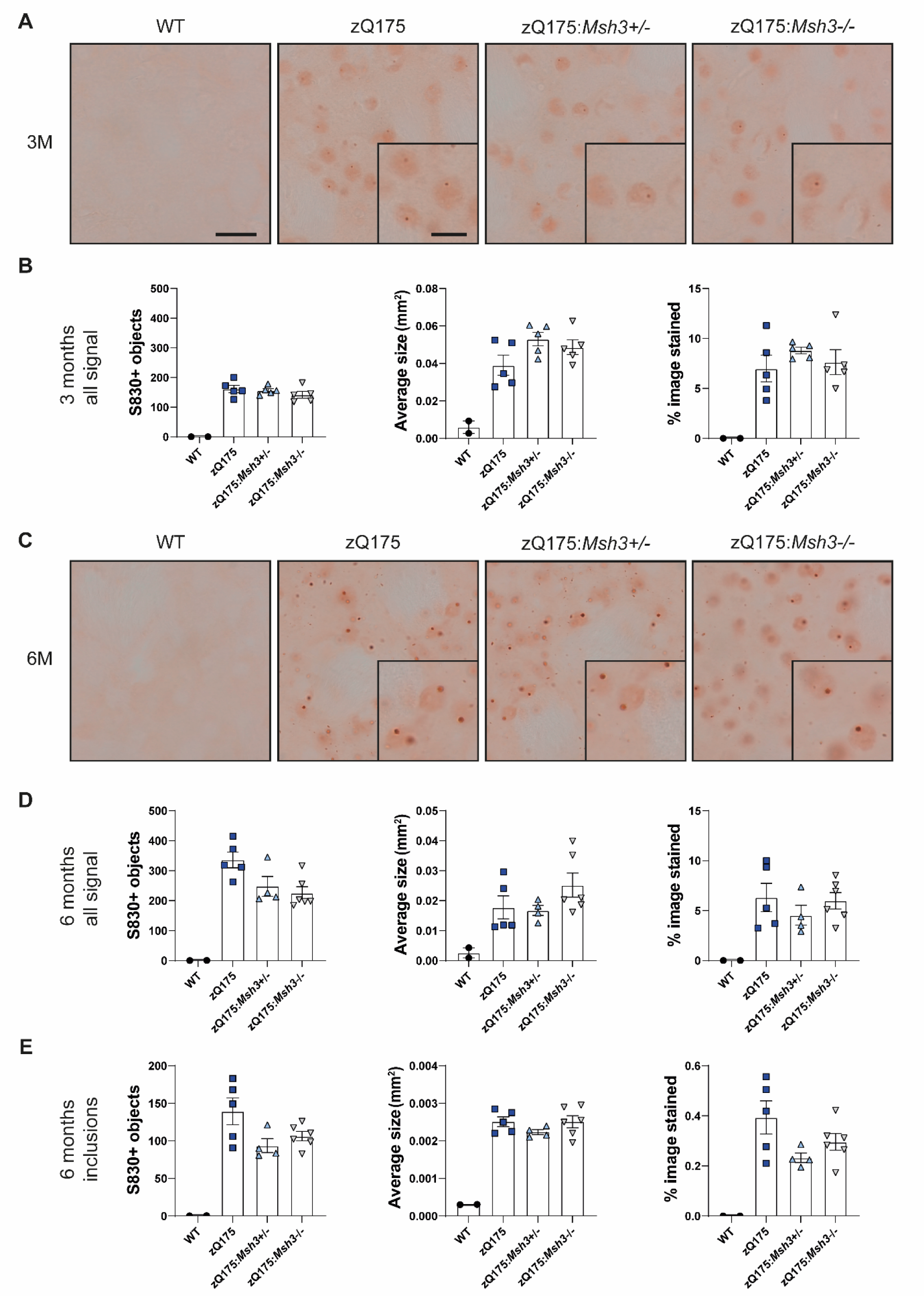
Striatal extranuclear, but not nuclear, HTT aggregation is reduced with loss of *Msh3*. Coronal striatal sections from WT, zQ175, zQ175:*Msh3*+/− and zQ175:*Msh3*-/- at 3 and 6 months of age were immunohistochemically stained with the S830 antibody. A nuclear counterstain was not applied to these images, as this would mask the nuclear signal. **(A)** HTT aggregation was detectable by 3 months of age in the zQ175 mice and S830 immunostaining levels were comparable irrespective of *Msh3* genotype. **(B)** Thresholding was applied to images to quantify levels of staining of S830 positive objects, average area of object, and the percentage of image covered. The *Msh3* genotype had no impact on these three measures of S830 staining. **(C)** At 6 months of age, in addition to diffuse nuclear aggregation, nuclear inclusions were also prominent in many nuclei. **(D-E)** Thresholding was applied to quantify the number of **(D)** S830 positive objects and **(E)** inclusions. In each case, the number of objects, the average area of the objects, and the percentage of an image covered by the S830 signal was determined. One-way ANOVA with Bonferroni correction. Error bars = SEM. *N* = 5/zQ175 genotype, *N* = 3 WT. Scale bar in large image = 10 μm, scale bar in insert = 5 μm. M = months, WT = wild-type.

We have previously established homogeneous time-resolved fluorescence (HTRF) assays to compare the levels of soluble and aggregated HTT isoforms between tissues.^43^ These included an assay specific for the soluble HTTexon1 protein (2B7-MW8) and a HTT aggregation assay (4C9-MW8) that we have shown is specific for the aggregated HTTexon1 protein.^7^ The epitope for 2B7 is within the first 17 amino acids of HTT, that for 4C9 is within the human-specific proline-rich region and MW8 acts as a C-terminal-neo-epitope antibody, and is specific for HTTexon1 in this HTRF assay (Fig. 7A). We applied these assays to the striatal, cortical and brain stem tissues of zQ175, zQ175:*Msh3*+/− and zQ175:*Msh3*-/- mice to investigate whether the prevention of somatic CAG repeat expansion had resulted in a decrease in HTT aggregation. Striatum, cortex, and brain stem from zQ175 and wild-type mice at two months of age were included in the analysis.

**Figure 7.**
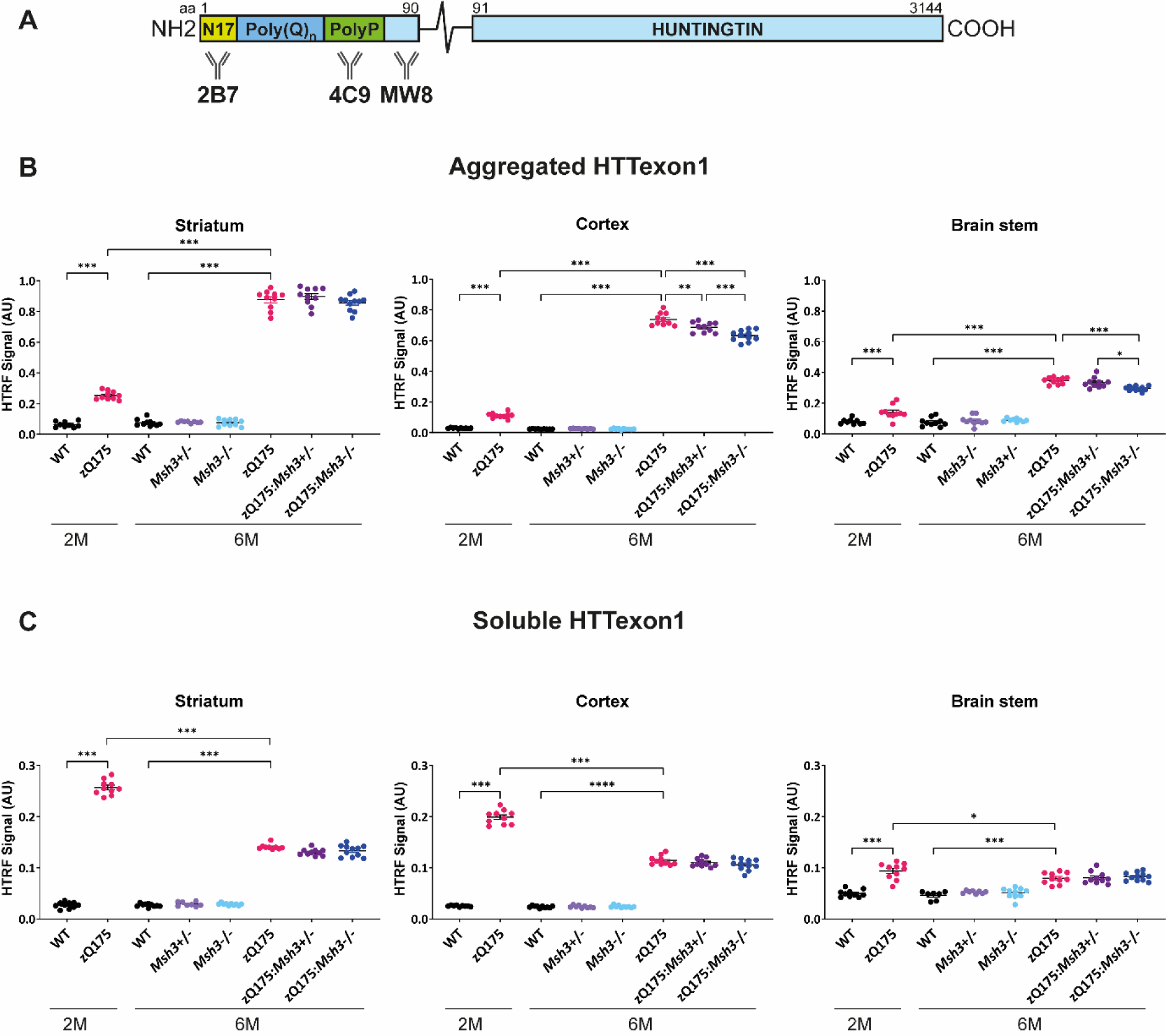
Striatal HTTexon1 aggregation is not affected by loss of *Msh3* expression. **(A)** Schematic of epitope location of antibodies used in HTTexon1 HTRF assays**. (B)** Levels of aggregated HTTexon1 as measured by the 4C9-MW8 assay indicated that aggregation increased from 2 to 6 months of age in zQ175 mice. There was no change in the levels of HTTexon1 aggregation detected between zQ175, zQ175:*Msh3*+/− and zQ175:*Msh3*+/− in the striatum, but small and significant differences were detected in the cortex and brainstem at 6 months of age. **(C)** High levels of soluble HTTexon1, as detected by the 2B7-MW8 assay, were present by 2 months of age in the zQ175 striatum, cortex and brain stem and had decreased at 6 months in all regions. There was no difference in soluble HTTexon1 at 6 months of age between the zQ175, zQ175:*Msh3*+/− or zQ175:*Msh3-*/- genotypes. 6-month samples: One-way ANOVA with Bonferroni correction; 2-month and 6-month zQ175 and WT: two-way ANOVA with Bonferroni correction. Error bars = SEM. \**P* ≤ 0.05, ** *P* ≤ 0.01, \*\*\**P* ≤ 0.001. *N* = 10/genotype. The test statistic, degrees of freedom and *p* values are summarised in Supplementary Table 5. AU= arbitrary units. WT= wild-type.

The 4C9-MW8 assay showed that HTTexon1 aggregation had increased from two to six months of age in all three brain regions (Fig. 7B). At six months of age, there was no difference in HTTexon1 aggregation between zQ175, zQ175:*Msh3+/−* and zQ175:*Msh3-/-* in the striatum, consistent with the immunohistochemical analysis (Fig. 7B). However, there was a small reduction HTTexon1 aggregation in the cortex of zQ175:*Msh3*+/− mice as compared to zQ175, and in zQ175:*Msh3*-/- mice as compared to zQ175:*Msh3*+/− (Fig. 7B). Similarly, the level of HTTexon1 aggregation in the brain stem of zQ175:*Msh3*-/- mice was decreased as compared to zQ175 and zQ175:*Msh3*+/− mice (Fig. 7B). The 2B7-MW8 HTRF assay demonstrated that the soluble levels of HTTexon1 in zQ175 mice remain unchanged by *Msh3* genotype, in all three brain regions (Fig. 7C). Consistent with this, the level of the *Htt1a* transcript was not consistently changed in the striatum, cortex and brain stem between zQ175, zQ175:*Msh3*+/− and zQ175:*Msh3*-/- (Supplementary Fig. 4).

## Discussion

Genetic variation in DNA mismatch repair pathway genes modifies the age of onset and rate of progression of Huntington’s disease by slowing the rate of somatic CAG repeat expansion. Of these, MSH3 has become a major focus for therapeutic development because loss of *MSH3* appears to be well tolerated.^36^ In preparation for the preclinical evaluation of these potential treatments, we genetically ablated *Msh3* in the zQ175 knock-in mouse model of Huntington’s disease to determine the maximum benefit of targeting *Msh3* in this model. Although loss of *Msh3* prevented somatic CAG repeat instability in striatal neurons, there was no effect on early molecular or pathological phenotypes. These data indicate that highly expanded CAG repeats can reach a threshold above which further expansion does not accelerate the onset of cellular pathology.

The zQ175 knock-in mouse model has been used extensively for mechanistic and preclinical studies, because its highly expanded CAG repeat results in molecular and pathological phenotypes at a younger age than in other knock-in mouse models of Huntington’s disease.^37,38^ However, somatic CAG repeat instability had not previously been characterised in these mice. We generated CAG-repeat traces for ten CNS regions and some peripheral tissues for zQ175 mice at six months of age, and compared these to traces from ear biopsies sampled when the mice were 12 days old. Pronounced instability was detected in the olfactory bulb, striatum, brain stem, colliculus, thalamus and spinal cord. This pattern of expansion was comparable to that previously published for the transgenic R6/1 mice with an repeat of (CAG)_115_,^49^ and so was not influenced by genomic context. In the periphery, there was considerable instability in the liver, consistent with that reported for the knock-in *Hdh*Q111 model.^23,33,49^ That the CAG repeat contracted in the pituitary was an unexpected finding, and could provide mechanistic insights in a future study. We have consistently found the heart CAG repeat to be comparatively stable and propose this could be used as a reference tissue, should an ear or tail biopsy from young mice not be available.

Capillary electrophoresis of bulk-PCR products, as used here, will not have captured the full extent of CAG repeat expansions for any brain region or tissue, as shorter CAG tracts will be preferentially detected. This is particularly challenging for the zQ175 model, for which the baseline CAG repeat length is already very high. This issue has been circumvented with small-pool PCR approaches whereby samples were diluted to contain a few DNA molecules and alleles were detected by Southern blotting.^4,52^ However, higher throughput analyses are required. A comparison of capillary electrophoresis, MiSeq or PacBio SMRT sequencing for bulk-tissue analysis demonstrated that PCR-based approaches cannot be used to estimate modal length or to quantify expansions greater than (CAG)_250_,^53^ and approaches that avoid an amplification step (no-amp) are under development.^54,55^ Bulk-PCR analysis also obscures any cell-type specificity for CAG repeat instability. Laser dissection showed that large expansions were present in neurons rather than glia in mouse models and *post-mortem* brains.^56,57^ More recently, CAG repeat expansions within a given cell population were determined by sorting cell-specific nuclei from Huntington’s disease *post-mortem* brains followed by high-throughput sequencing. This showed that, in the striatum, expansions had occurred in medium spiny neurons and CHAT+ interneurons but not in glial populations.^58^ Interestingly, CAG repeat expansions were detected in cerebellar Purkinje cells, a rare cell type in this brain structure,^58^ whereas CAG repeat instability in the cerebellum had not been uncovered by small-pool PCR.^4^

The ablation of MSH3 completely prevented somatic CAG repeat expansion in all brain regions and peripheral tissues and in many cases resulted in a reduction of the repeat length mode by up to five CAGs. Comparison of the instability index for zQ175, zQ175:*Msh3*+/− and zQ175:*Msh3*-/- demonstrated that for most brain regions and tissues, a 50% reduction in MSH3 resulted in a 50% decrease in CAG repeat expansion. In the case of the striatum and spinal cord, the reduction in expansion between zQ175 and zQ175:*Msh3*+/− mice was much less than 50%, possibly reflecting an underestimate of the extent of expansion in these brain regions, as discussed above. MSH3 levels only affected expansion, as neither a loss of MSH3, or a reduction in its levels reversed the CAG repeat contraction observed in the pituitary.

Our rationale for performing the intercross between zQ175 and *Msh3* knock-out mice was to determine whether zQ175 might be useful preclinically for assessing therapeutics targeting MSH3. At six months of age zQ175 mice have robust molecular and pathological phenotypes in the form of striatal transcriptional dysregulation^13^ and the deposition of aggregated HTT throughout the brain,^7^ and have been used in a zQ175 preclinical trial to assess HTT-lowering with zinc finger proteins.^39^ Therefore, we performed RNA-seq on striatal RNA for wild-type, *Msh3*-/-, zQ175 and zQ175:*Msh3*-/- mice. Remarkably, the ablation of MSH3 had no impact on the dysregulated transcriptional profile of the zQ175 striatum; of 2,486 dysregulated genes, only 3% of dysregulated genes had an expression that was reversed toward wild-type levels, and in 1%, this was exacerbated; changes that would be below background levels. This is in stark contrast to the effect of ablating MSH3 in CAG140 knock-in mice (zQ175 mice are CAG140 mice with a longer CAG repeat) in which HTT aggregation in striatal nuclei was decreased and a considerable percentage of the dysregulated transcripts were reversed toward wild-type levels (X. William Yang, personal communication, manuscript submitted).

It is now widely accepted that it is the somatic CAG repeat expansion in the brain that drives the age of onset and rate of progression of Huntington’s disease,^59^ but the mechanism through which this initiates pathogenesis has not been confirmed. The alternative processing of the *HTT* pre-mRNA to generate *HTT1a*, provides a candidate for this second step.^5,6^ The production of *HTT1a* increases with increasing CAG repeat length and it encodes the HTTexon1 protein that is known to be highly aggregation-prone and pathogenic.^15^ Therefore somatic CAG repeat expansion would be expected to increase the levels of *Htt1a* and of the soluble HTTexon1 protein at the single cell level. However, when comparing zQ175:*Msh3*-/- mice with zQ175, we did not identify a reduction in *Htt1a* or HTTexon1 in bulk tissue by QuantiGene or HTRF analyses, respectively. These techniques may not be sufficiently sensitive to detect these changes at the tissue-level, given that the CAG repeat in zQ175 mice is already highly expanded and that in most cells further expansions do not increase the CAGs repeat length to a dramatic degree.

Changes in HTT aggregation might be more amenable to detection, given the cumulative nature of the process and that only the cells in which aggregation occurs contribute to the analysis. HTT aggregation is a concentration-dependent process^60^ and whether this occurs in a given subcellular location will depend on a number of factors e.g. the concentration of HTTexon1 and other HTT N-terminal fragments, the polyQ length (as the longer the polyQ tract the lower the concentration required), post-translational modifications, interacting proteins or RNAs, and the proteostasis network. In zQ175 mice, it is the soluble HTTexon1 protein that is depleted from brain regions during the aggregation process.^7^ The N-terminus of HTT contains a potent nuclear export signal,^61,62^ and HTT is retained in neuronal nuclei because it aggregates there, appearing first as a diffuse immunoreactive product that fills the nucleus.^15,61,63^ The threshold concentration for the nucleation of aggregation will be cell-type and subcellular location specific.

Knocking out the mismatch repair genes *Msh2* or *Mlh1* delayed the accumulation of aggregated HTT in striatal nuclei of the *Hdh*Q111 knock-in model.^22,23^ However, prevention of somatic CAG repeat expansion in zQ175 mice had no effect on the deposition of aggregated HTT in striatal nuclei. Given that HTT aggregation precedes transcriptional changes,^7,13,15^ this was consistent with MSH3 ablation having no effect on transcriptional dysregulation in the zQ175 striatum. Therefore, in this subcellular location, (CAG)_185_ in the *Htt* gene produced HTTexon1 at or above the threshold concentration needed to initiate aggregation, and further CAG expansion could not accelerate this process. It is striking that highly expanded CAG repeats (of greater than (CAG)_200_) in humans cause onset of Huntington’s disease before two years of age, whereas those of (CAG)_100–130_ caused onset at around four years.^64,65^ This suggests that for CAG repeats approaching (CAG)_200_, initiation of the pathogenic process does not require further expansion through somatic CAG repeat instability.

MSH3 has significant therapeutic potential for Huntington’s disease and other trinucleotide repeat disorders,^34^ and a wide range of MSH3-targeting modalities are under development.^29,66^ Our data indicate that whilst ablation of MSH3 prevented somatic CAG repeat expansion in zQ175 mice, this model will not be useful for evaluating these MSH3-targeting treatments, and that knock-in mice with shorter CAG repeat mutations should be used. However, more importantly, we found that somatic expansions in zQ175 striatal neurons did not accelerate HTT aggregate pathology or transcriptional dysregulation, indicative of a pathogenic repeat threshold at, or close to, (CAG)_185_. Early childhood onset cases of Huntington’s disease with repeat expansions close to (CAG)_200_, are consistent with a pathogenic threshold occurring at around this repeat length. Available data indicate that successful MSH3-targeting treatments will slow down the rate of somatic CAG repeat expansion in the brains of Huntington’s disease mutation carriers. The prevention of somatic CAG repeat expansion is unlikely as that would require MSH3 ablation. The trajectory for repeat expansion in the Huntington’s disease brain is not understood and these results indicate that the greatest benefit for MSH3-targeting therapeutics will be realised by treating as early as possible before cell-specific pathogenic thresholds have been achieved.

## Abbreviations

CRISPR: clustered regularly interspaced short palindromic repeats
DAB: 3.3′-diaminobenzidine
FELASA: Federation of European Laboratory Animal Science Associations
gRNA: guide RNA
GOBP: gene ontology biological process
GWAS: genome-wide association study
HTRF: homogeneous time-resolved fluorescence
HTTexon1: exon 1 HTT protein
IQR: interquartile range
PBS: phosphate buffer saline
PCA: principal component analysis
polyA: polyadenylation
polyQ: polyglutamine
Q: glutamine
qPCR: real-time quantitative PCR
RNA-seq: RNA sequencing
SEM: standard error of the mean
SMRT: single molecule real-time sequencing
WT: wild-type.

## Acknowledgements

The authors wish to thank Jian Chen at the CHDI Foundation for advice on RNA sequencing and facilitating the data analysis. We also thank Kirupa Sathasivam and Pamela Farshim for help with sectioning brains and Casandra Gomez-Paredes for help with QuantiGene analysis.

## Funding

This work was supported by grants from the CHDI Foundation, UK Dementia Research Institute, which receives its funding from Dementia Research Institute Ltd, funded by the Medical Research Council, Alzheimer’s Society and Alzheimer’s Research UK and the Wellcome Trust (223082/Z/21/Z). Mice were generated with support from NIH grants R01CA248536 and 5P30CA13330.

## Competing interests

JRG and MBH are employees of Rancho BioSciences and CLB is an employee of LoQus23 Therapeutics.

## Data availability

RNA-seq data that support the finding of this study have been deposited in GEO with the accession no. GSE245510. The authors confirm that all other data supporting the findings in this study are available within the article and its Supplementary Material. Raw data will be shared by the corresponding author upon request.

