## Supplementary material for "A CAG repeat threshold for therapeutics targeting somatic instability in Huntington’s disease"

**SUPPLEMETARY MATERIAL**

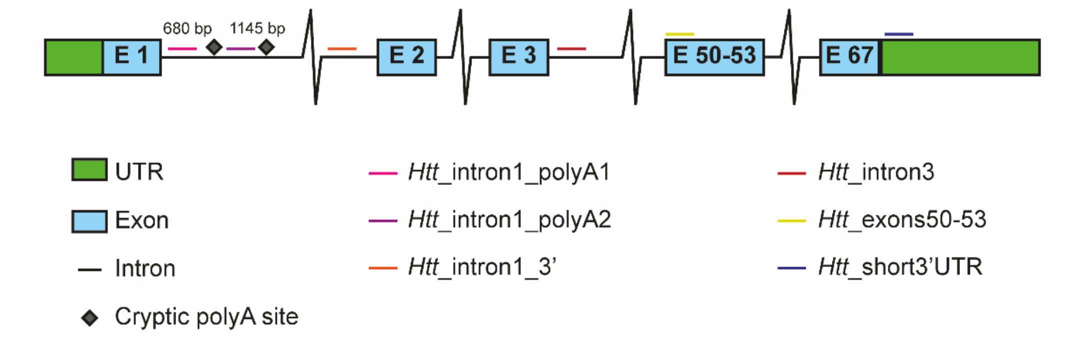

**Supplementary Figure 1 Schematic showing the position of the QuantiGene probes across the *Htt* gene.** The *Htt*_intron1_polyA1 and *Htt*_intron1_polyA2 probes are located before the activated polyA sites and detect the *Htt1a* transcript. The Htt_intron1_3’ and Htt_intron3 probes detect unspliced pre-mRNA. The Htt_exons50-53 and Htt_short3’UTR probes detect the full-length processed mRNA. UTR = untranslated region, bp = base pairs.

**Supplementary Table 1. The multiplex QuantiGene assay for the detection of *Htt* transcripts.**

| ***Htt* plex** | **Accession number** | **Specific location** | **Target** |
|---|---|---|---|
| *Htt*_intron1_pA1 | GS03082 | 521-983 | |
| *Htt*_intron1_pA2 | GS03084 | 1104-1465 | |
| *Htt*_intron1_3’ | GS03085 | 16339-16922 | |
| *Htt*_intron3 | GS03083 | 30195-30846 | Huntingtin |
| *Htt*_exons 50-53 | NM_010414 | 6901-7433 | |
| *Htt*_short 3’UTR | NM_010414 | 9553-9993 | |
| *Eif4a2* | NM_013506 | 710-1271 | |
| *Rpl13a* | NM_009438 | 2-467 | Housekeeping gene |
| *Canx* | NM_007597 | 1195-1720 | |
| *Atp5b* | NM_016774 | 22-406 | |

**Supplementary Table 2. The multiplex QuantiGene assay for the detection of DNA mis-match repair transcripts.**

| ***MMR* plex** | **Accession number** | **Specific location** | **Target** |
|---|---|---|---|
| *Msh3 4_7* | NM_010829.2 | 582-1140 | |
| *Msh2* | NM_008628.3 | 2260-2727 | |
| *Msh6* | NM_010830.2 | 3638-4124 | |
| *Exo1** | NM_012012.4 | 4041-4496 | Gene of interest |
| *Mlh1* | NM_026810.2 | 1715-2065 | |
| *Mlh3* | NM_175337.2 | 3393-3864 | |
| *Pms1* | NM_153556.2 | 2464-2878 | |
| *Pms2* | NM_008886.2 | 1660-2043 | |
| *Fan1* | NM_177893.4 | 2437-2585 | |
| *Eif4a2* | NM_013506 | 710-1271 | |
| *Rpl13a* | NM_009438 | 2-467 | Housekeeping gene |
| *Canx* | NM_007597 | 1195-1720 | |
| *Atp5b* | NM_016774 | 22-406 | |

**Exo1* not detected above background in any region analysed.

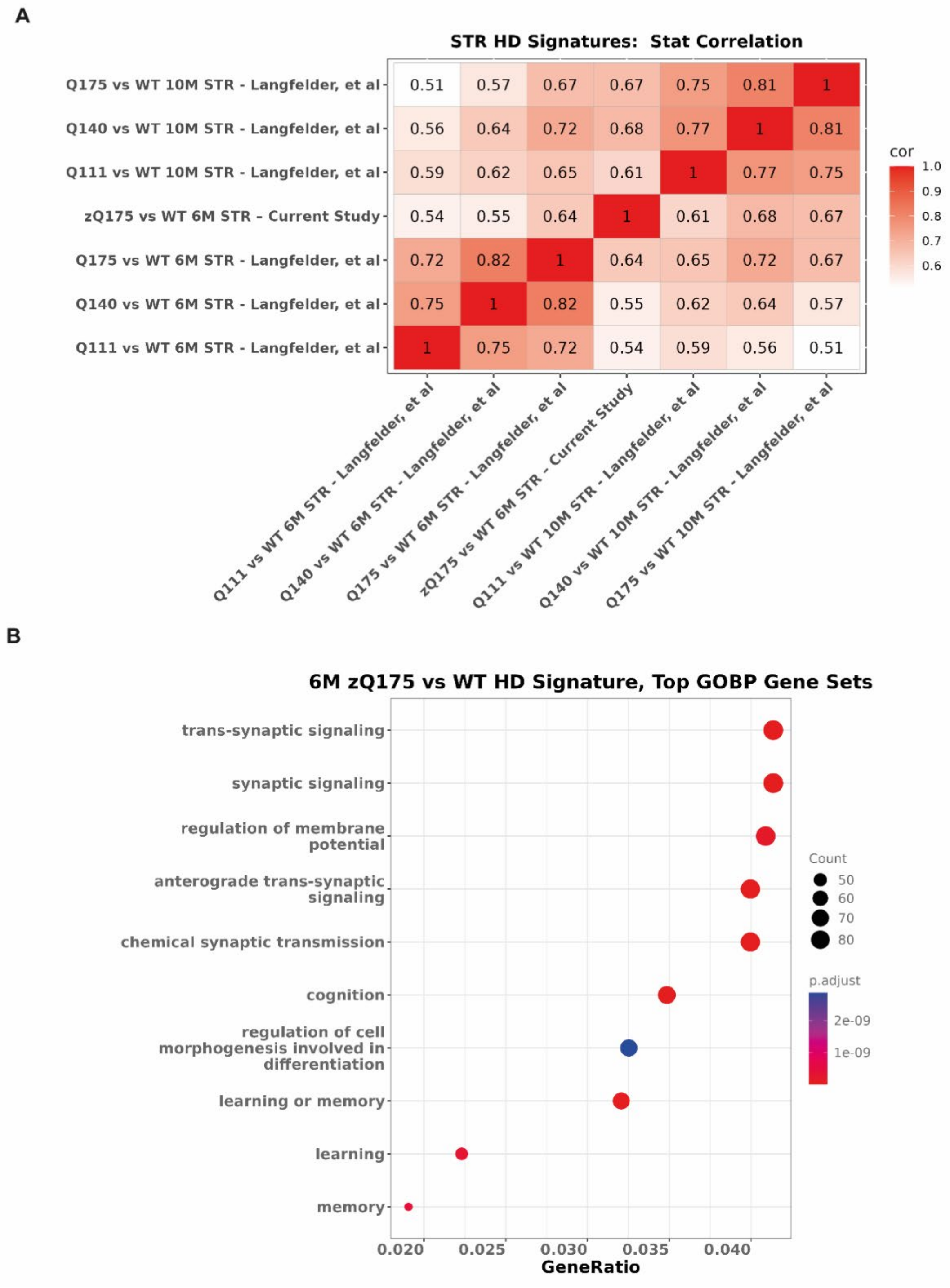

**Supplementary Figure 2. Comparison of the transcriptional dysregulation signature in zQ175 mice with that in knock-in models published previously. (A)** Heatmap for dysregulated genes in the zQ175 striatum in this 'current study' at 6 months of age compared to dysregulated data sets, for zQ175, Q140 and Q111 knock-in models at 6 and 10 months of age, using data from Langfelder et al. (2016).[1] There is significant overlap in the dysregulation detected in the current study with that in the previous datasets. **(B)** Gene set overrepresentation of the total 2,486 genes dysregulated in the zQ175 striatum analysed against the GOBP to identify biological pathways that are dysregulated. Of the 227gene sets that had $q < 0.01$, the top 10 ranked by the GeneRatio, which is the number of dysregulated genes in the given gene set divided by the number of dysregulated genes present overall in the GOBP (gene ontology biological process) gene set collection.

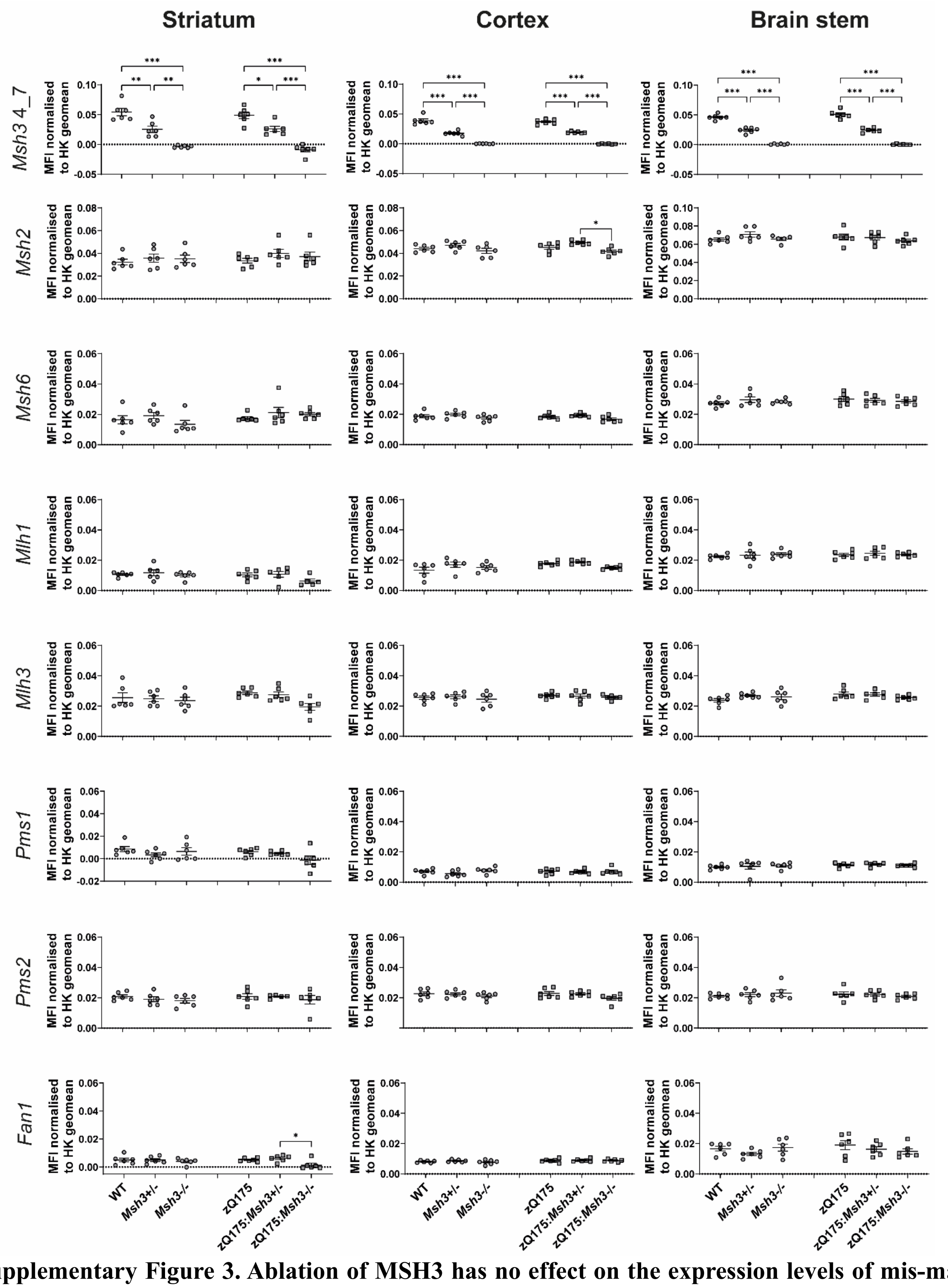

**Supplementary Figure 3. Ablation of MSH3 has no effect on the expression levels of mis-match repair gene transcripts.** QuantiGene analysis of the levels of DNA mis-match repair transcripts measured in cortex, striatum, and brainstem. The *Msh34_7* assay is located in deleted region of the genetically modified *Msh3* gene and levels were reduced in *Msh3*+/- and zQ175:*Msh3*+/- mice and ablated in *Msh3*+/- and zQ175:*Msh3*+/- mice, as expected. The other mis-match repair transcripts show no consistent change in expression level. One-way ANOVA with Tukey's correction. Error bars = SEM. *$P \leq 0.05$, ** $P \leq 0.01$, ***$P \leq 0.001$. $N = 6$ /genotype. The test statistic, degrees of freedom and *p* values are summarised in **Supplementary Table 6.** WT = wild-type, MFI = mean fluorescence intensity HK = housekeeping genes.

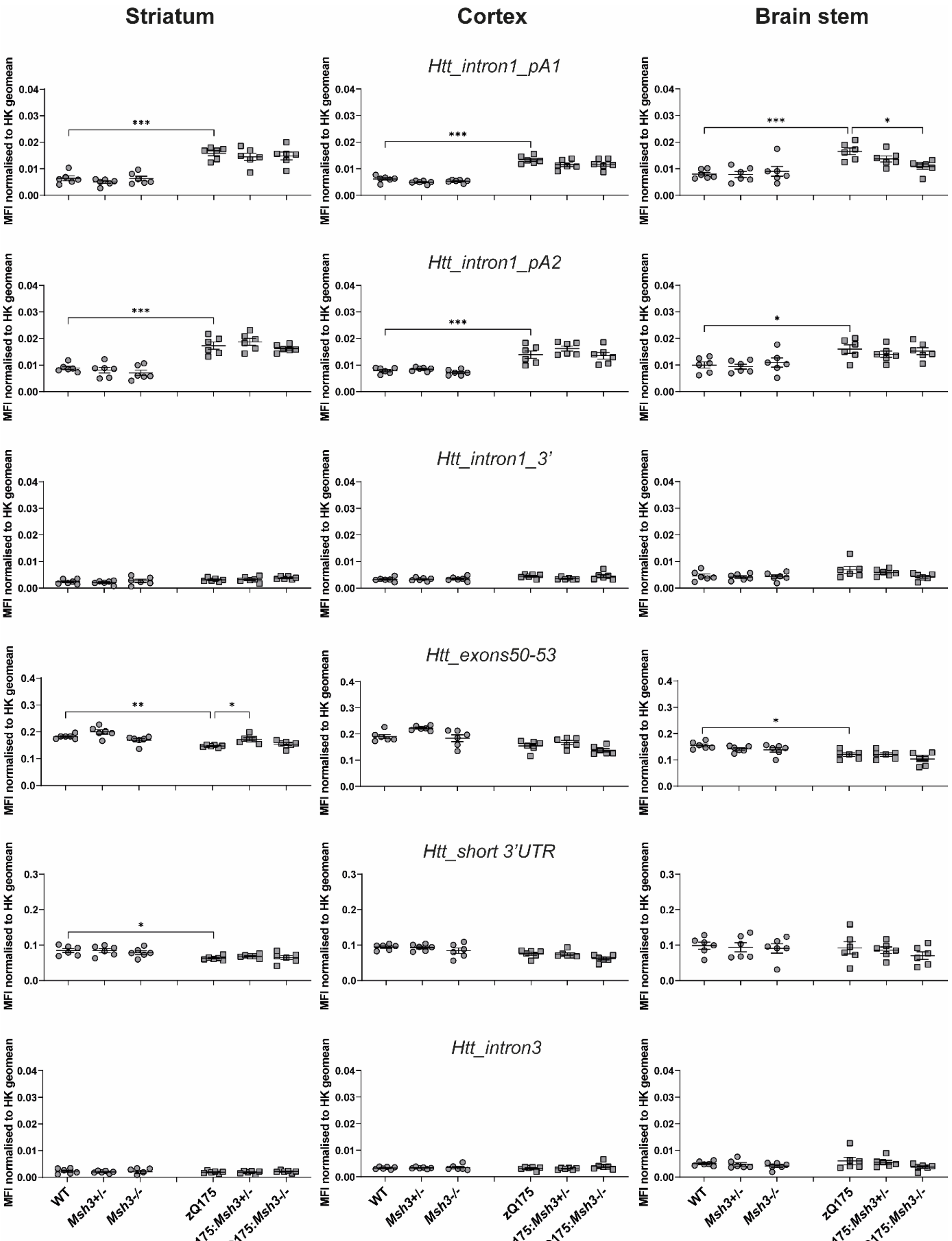

**Supplementary Figure 4 Ablation of MSH3 has no effect on the expression levels or processing of *Htt* transcripts.** QuantiGene analysis of full-length *Htt* and *Htt1a* transcripts in the striatum, cortex, and brainstem. *Htt1a* was detected in zQ175 mice at 6 months of age by *Htt*_intron1_polyA1 and *Htt*_intron1_polyA2 assays. The *Msh3* genotype had no consistent effect on *Htt1a* levels or full-length *Htt* levels. The *Htt*_intron1_3' and *Htt*_intron3 probes acted as pre-mRNA controls. One-way ANOVA with Tukey's correction. Error bars = SEM. *$P \leq 0.05$, ** $P \leq 0.01$, ***$P \leq 0.001$. $N = 6$ /genotype. The test statistic, degrees of freedom and $p$ values are summarised in **Supplementary Table 7.** WT = wild-type, MFI = mean fluorescence intensity HK = housekeeping genes.

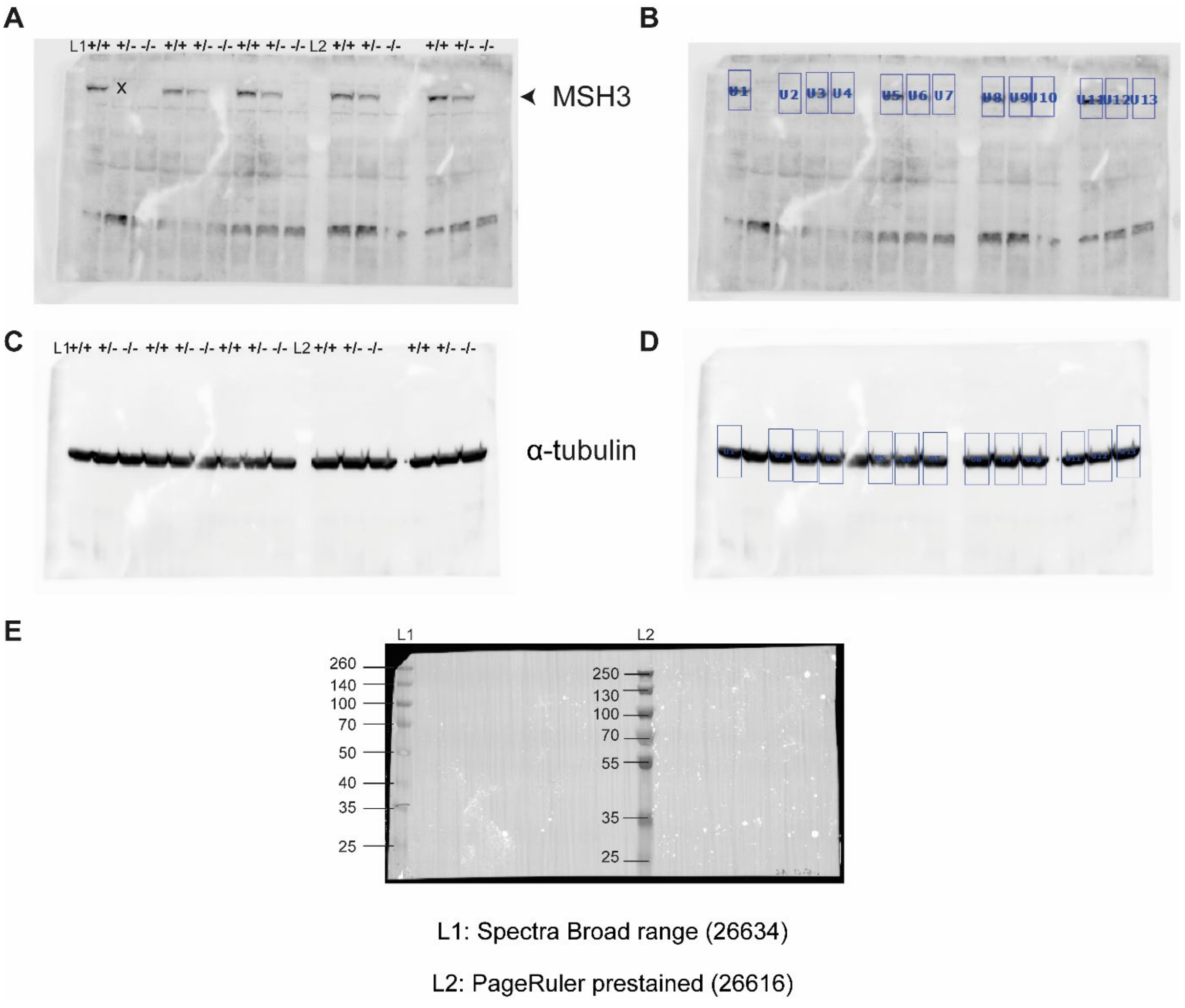

**Supplementary Figure 5. Full-length western blots for Figure 1C. (A)** membrane was probed with the MSH3 antibody and the expected band of ~130 kDa was detected. X indicates likely bubble/transfer issue. **(B)** MSH3 band intensity was quantified using BioRad ImageLab volume tool, using rectangles of same size across all lanes. (**C**) Blot was then probed with the α-tubulin antibody and **(D)** quantified with ImageLab. (**E**) Image of molecular weight sizing ladders and band sizes.

**Supplementary Table 3. The test statistic, degrees of freedom and *p* values for the one-way ANOVA of data presented in Figure 1B and 1C**

| Cortex, Msh3 quantification | | |
|---|---|---|
| RNA | F (2, 13) = 32.03 | P<0.001 |
| Protein | F (2, 10) = 15.29 | P<0.001 |

**Supplementary Table 4. The mixed effects analysis of data presented in Figure 3A and 3B.**

| | **Instability index** | | **Change in mode** | |
|---|---|---|---|---|
| Row Factor (Tissue) | F(5.1, 103.0) = 291.2 | P<0.0001 | F(5.9, 117.4) = 270.6 | P<0.0001 |
| Column Factor (Genotype) | F(2, 21) = 375.1 | P<0.0001 | F(2, 21) = 103.2 | P<0.0001 |
| Row Factor (Tissue x Genotype) | F(26, 264) = 32.02 | P<0.0001 | F(26, 260) = 18.63 | P<0.0001 |
| Random effects | SD | Variance | SD | Variance |
| Subject (Mouse) | 0.3914 | 0.1532 | 0.4406 | 0.1942 |
| Residual | 0.8791 | 0.7728 | 0.8198 | 0.6721 |

SD = standard deviation

**Supplementary Table 5. The test statistic, degrees of freedom and *p* values for the data presented in Figure 7**

Two-way ANOVA for the comparison of wild-type and zQ175 mice at 2 and 6 months of age.

| | **4C9– MW8** | | | | | |
|---|---|---|---|---|---|---|
| | Striatum | | Cortex | | Brain stem | |
| Age*Genotype | F (1, 35) = 635.9 | P<0.001 | F (1, 35) = 1893 | P<0.001 | F (1, 35) = 121.6 | P<0.001 |
| Age | F (1, 35) = 682.9 | P<0.001 | F (1, 35) = 1825 | P<0.001 | F (1, 35) = 115.4 | P<0.001 |
| Genotype | F (1, 35) = 1663 | P<0.001 | F (1, 35) = 3000 | P<0.001 | F (1, 35) = 305.7 | P<0.001 |
| | **2B7– MW8** | | | | | |
| | Striatum | | Cortex | | Brain stem | |
| Age*Genotype | F (1, 35) = 462.0 | P<0.001 | F (1, 35) = 232.4 | P<0.001 | F (1, 33) = 1.928 | P<0.001 |
| Age | F (1, 35) = 462.0 | P<0.001 | F (1, 35) = 251.0 | P<0.001 | F (1, 33) = 5.730 | P<0.001 |
| Genotype | F (1, 35) = 4046 | P<0.001 | F (1, 35) = 2345 | P<0.001 | F (1, 33) = 104.5 | P<0.001 |

One-way ANOVA for the comparison of the six genotypes at 6 months of age.

| **4C9– MW8** | | | | | |
|---|---|---|---|---|---|
| Striatum | | Cortex | | Brain stem | |
| F (5, 54) = 295.3 | P<0.001 | F (5, 53) = 1805 | P<0.001 | F (5, 54) = 295.3 | P<0.001 |
| **2B7– MW8** | | | | | |
| Striatum | | Cortex | | Brain stem | |
| *H* =45.1, df =57 | P<0.001 | F (5, 54) = 462.8 | P<0.001 | F (5, 50) = 30.75 | P<0.001 |

**Supplementary Table 6. The test statistic, degrees of freedom and *p* values for the one-way ANOVA presented in Supplementary Figure 3**

| | Cortex | | Striatum | | Brain stem | |
|---|---|---|---|---|---|---|
| ***Msh3 4_7*** | F (5, 30) = 124.8 | P<0.001 | F (5, 30) = 31.98 | P<0.001 | F (5, 30) = 164.2 | P<0.001 |
| ***Msh2*** | F (5, 30) = 3.751 | P=0.009 | F (5, 30) = 0.7020 | P=0.63 | F (5, 29) = 0.9925 | P=0.44 |
| ***Msh6*** | F (5, 30) = 1.994 | P=0.11 | F (5, 30) = 1.421 | P=0.25 | F (5, 30) = 0.5684 | P=0.72 |
| ***Mlh1*** | F (5, 30) = 2.580 | P=0.05 | F (5, 29) = 1.586 | P=0.20 | F (5, 30) = 0.3694 | P=0.87 |
| ***Mlh3*** | F (5, 30) = 0.4578 | P=0.80 | F (5, 30) = 2.219 | P=0.08 | F (5, 30) = 1.612 | P=0.19 |
| ***Pms1*** | F (5, 30) = 0.9195 | P=0.48 | F (5, 30) = 1.975 | P=0.11 | F (5, 30) = 0.4686 | P=0.80 |
| ***Pms2*** | F (5, 30) = 1.359 | P=0.27 | F (5, 29) = 0.5190 | P=0.76 | F (5, 30) = 0.3698 | P=0.87 |
| ***Fan1*** | F (5, 30) = 1.202 | P=0.33 | F (5, 30) = 2.497 | P=0.05 | F (5, 30) = 1.017 | P=0.43 |

**Supplementary Table 7. The test statistic, degrees of freedom and *p* values for the one-way ANOVA presented in Supplementary Figure 4**

| Assay | Cortex | | Striatum | | Brain stem | |
|---|---|---|---|---|---|---|
| *Htt*_intron1_pA1 | F(5, 30) = 17.55 | P<0.0001 | F (5, 30) = 23.04 | P<0.0001 | F (5, 30) = 8.143 | P<0.0001 |
| *Htt*_intron1_pA2 | F (5, 30) = 47.24 | P<0.0001 | F (5, 30) = 25.17 | P<0.0001 | F (5, 30) = 4.575 | P=0.0032 |
| *Htt*_intron1_3' | F (5, 30) = 2.613 | P=0.0446 | F (5, 30) = 2.649 | P=0.0425 | F (5, 30) = 2.208 | P=0.0796 |
| *Htt*_intron3 | F (5, 30) = 12.43 | P<0.0001 | F (5, 30) = 11.14 | P<0.0001 | F (5, 30) = 6.694 | P=0.0003 |
| *Htt*_exons50_53 | F (5, 30) = 6.632 | P=0.0003 | F (5, 30) = 4.240 | P=0.0049 | F (5, 30) = 0.6347 | P=0.6748 |
| *Htt*_short3'UTR | F (5, 30) = 1.094 | P=0.3842 | F (5, 30) = 0.2239 | P=0.9493 | F (5, 30) = 1.233 | P=0.3185 |
